# A novel qPCR assay to detect the presence of *Anopheles gambiae* complex mosquitoes

**DOI:** 10.64898/2026.03.03.707393

**Authors:** David Richard Hemprich-Bennett, Gonçalo Alves, Abigail Bailey, Fred Aboagye-Antwi, Owen T. Lewis, Talya D. Hackett

## Abstract

**Background:** *Anopheles* mosquitoes vector pathogens responsible for more than 600,000 human deaths annually. Ecological studies of these insects are important to guide effective vector-control campaigns and to understand their broader ecological consequences. Molecular ecology methods, particularly qPCR, provide a valuable tool in such studies. By detecting trace DNA of a taxon of interest within mixed or environmental samples, qPCR can facilitate identification of prey taxa of interest in the diets of consumers. However, no protocol for the detection of *An. gambiae* complex mosquitoes in dietary samples has been available.

**Methods:** We introduce a new set of qPCR primers (Agam_CO1_F1 and Agam_CO1_R1) and a probe-based assay for detection of *Anopheles gambiae*-complex mosquitoes, even with short reads common in dietary and environmental samples. The primers were tested *in vitro* for their specificity and sensitivity, and *in silico* using Primer-BLAST to assess potential off-target amplification.

**Results:** The qPCR primers amplified *An. gambiae* DNA even at low starting concentrations (5 copies µl^*-*1^). The primers did not amplify any non-target DNA in either the *in vitro* or *in silico* tests, but consistently amplified *An. gambiae* complex DNA. The primers can therefore provide reliable tests for the presence or absence of *An. gambiae* complex in dietary or eDNA samples.

**Conclusions:** The new qPCR primers should allow advances in research into mosquito ecology by allowing detection of even trace amounts of *An. gambiae* DNA in dietary and environmental samples.

**Graphical abstract:** 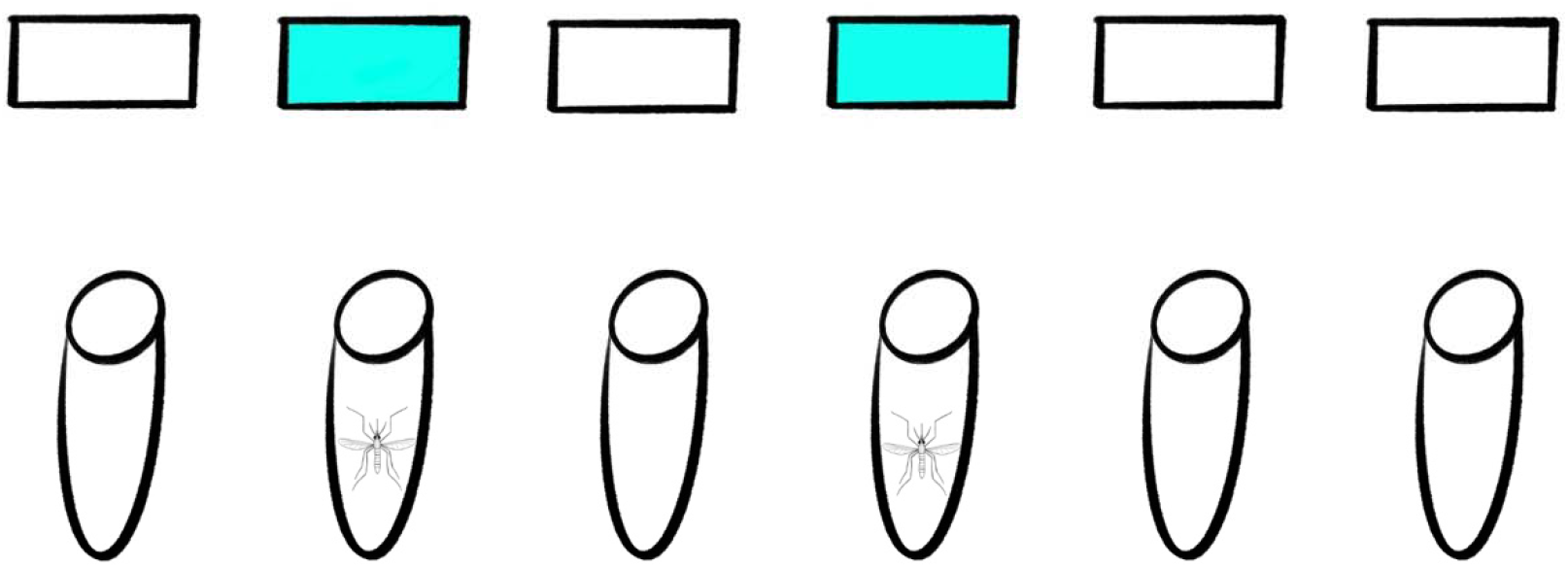

## Introduction

Malaria, the disease caused by parasites in the genus *Plasmodium*, remains a significant global disease burden for humans, with an estimated 282 million cases and 610,000 deaths in 2024, mostly children under 5 [1]. While a variety of *Plasmodium* species are vectored by a range of mosquito taxa to multiple host species, more than 90% of human malaria mortality is from *P. falciparum*, transmitted by mosquitoes in the genus *Anopheles* [2]. Malaria control campaigns in Africa frequently focus on the *Anopheles gambiae* complex, comprising the closely related species *An. amharicus, An. arabiensis, An. bwambae, An. coluzzii, An. gambiae sensu stricto* (s.s.), *An. melas, An. merus, An. quadriannulatus* [3]. Of these, *An. gambiae* s.s., *An. coluzzii* and *An. arabiensis* are the key human malaria vectors and remain genetically closely related, with evidence of limited hybridisation and introgression in the field [4–7].

A common approach to control malaria is to target the key mosquito vectors in the *Anopheles gambiae* complex. Many traditional vector control methods, including insecticide treated bed nets, indoor residual spraying and larvicide application, as well as proposed technologies such as population-suppression genetically modified mosquitoes, seek to reduce abundances and population sizes of *An. gambiae* [8]. Yet, we still have limited understanding of the broader ecological role *An. gambiae* plays in the wider ecosystem; an important consideration for predicting the likely consequences of their suppression [9]. The distributions, demography and ecological roles of species are strongly influenced by their interactions with their biotic and abiotic environment. There is some evidence for predation of adult *An. gambiae* by a spider species within the Salticidae [10], and many taxa are known or thought to feed on their larvae [11–13], but there is no evidence of strong dietary dependence on them by any species. Predicting the efficacy of malaria control-interventions and future range expansions relies on deepening this ecological knowledge and establishing, among other things, the trophic interactions of *An. gambiae* and insectivorous predators which might be feeding on them [13].

Traditionally, study of predatory interactions required either directly observing feeding interactions or visually identifying animal body parts in a predator’s faeces [14] or regurgitate [15]. Both methods are time and labour intensive, miss cryptic interactions and lack species specificity. More recently, advances in qPCR and DNA metabarcoding have made it possible to identify the DNA of prey items reliably and affordably within a predator or its excreta [16,17].

DNA metabarcoding, allowing the detection of multiple species in mixed samples (e.g. faeces, eDNA, pooled collections), is an established technique for the analysis of cryptic predation [18– 20]. However, primer biases and PCR stochasticity result in weaknesses when a given prey species is of particular interest, or when a predator consumes a wide variety of prey species [21,22]. PCR is inherently stochastic, and the use of generalist primers comes with the compromise that some species present may go undetected by chance, while others are over-amplified relative to their abundance [21,22]. The absence of a detection is therefore not evidence that a species is absent from a sample. Conventional PCR and qPCR can offer an alternative approach for dietary analysis, using primers which are designed to amplify only a narrow taxonomic breadth of potential prey, typically just a single species [17,23]. In conventional PCR, the simple presence or absence of a band in gel electrophoresis is used to determine the presence of a prey taxon’s DNA in a sample. qPCR provides additional information, using fluorescent compounds which are detected during successful amplification of DNA. This detection of fluorescing DNA allows PCR reactions to be monitored in real time. This provides data which can be used to compare the abundance of template DNA among samples [17], although this requires extensive calibration [24]. qPCR can thus be used to confirm the presence or absence of a specific prey taxon that researchers may have *a priori* expectation of or a specific interest in.

Given the inherent stochasticity in metabarcoding detection of any dietary taxa of interest and the specific nature of qPCR, it can be valuable to combine both DNA metabarcoding and qPCR when studying the predation of a particular species [17]. Metabarcoding can provide a broad overview of the diet of a predator, and qPCR can then validate the presence/absence of specific target prey taxa in a sample. While molecular methods have long been used to distinguish *Anopheles* species [25–27], most rely on amplicons that are prohibitively long for degraded DNA typical in dietary or eDNA analyses. Where assays have been developed for the detection of *An. gambiae*, they have either been designed solely to amplify individual subspecies within the complex and are longer than suitable for the analysis of dietary DNA [28], or amplified genes which are relatively understudied [29], reducing our ability to assess the possibility of non-target amplifications.

Here we present and validate qPCR primers targeting the CO1 gene (the gene most commonly used for DNA barcoding) developed specifically to detect *Anopheles gambiae* complex mosquitoes in predator diet. Many vector control campaigns affect the whole complex [9,30], therefore, we aimed to detect the presence of species from the complex as a whole, rather than individual species within it.

## Methods

### qPCR primer design

We used Geneious Prime 2023.2.1 (https://www.geneious.com) to design initial candidate qPCR primers. Culicid mitochondrial genomes or CO1 genes were downloaded from NCBI, and then aligned using the MAFFT algorithm [31,32]. We then visually searched for regions of similarity within the *An. gambiae* complex that were distinct from those in other members of the *Anopheles* and other culicid genera. The primers were further refined against an alignment of the *An. gambiae* complex in Bioedit 5.9.0, and a probe-based qPCR assay was developed, as shown in Table 1. The resulting amplicon is 192bp.

**Table 1:**
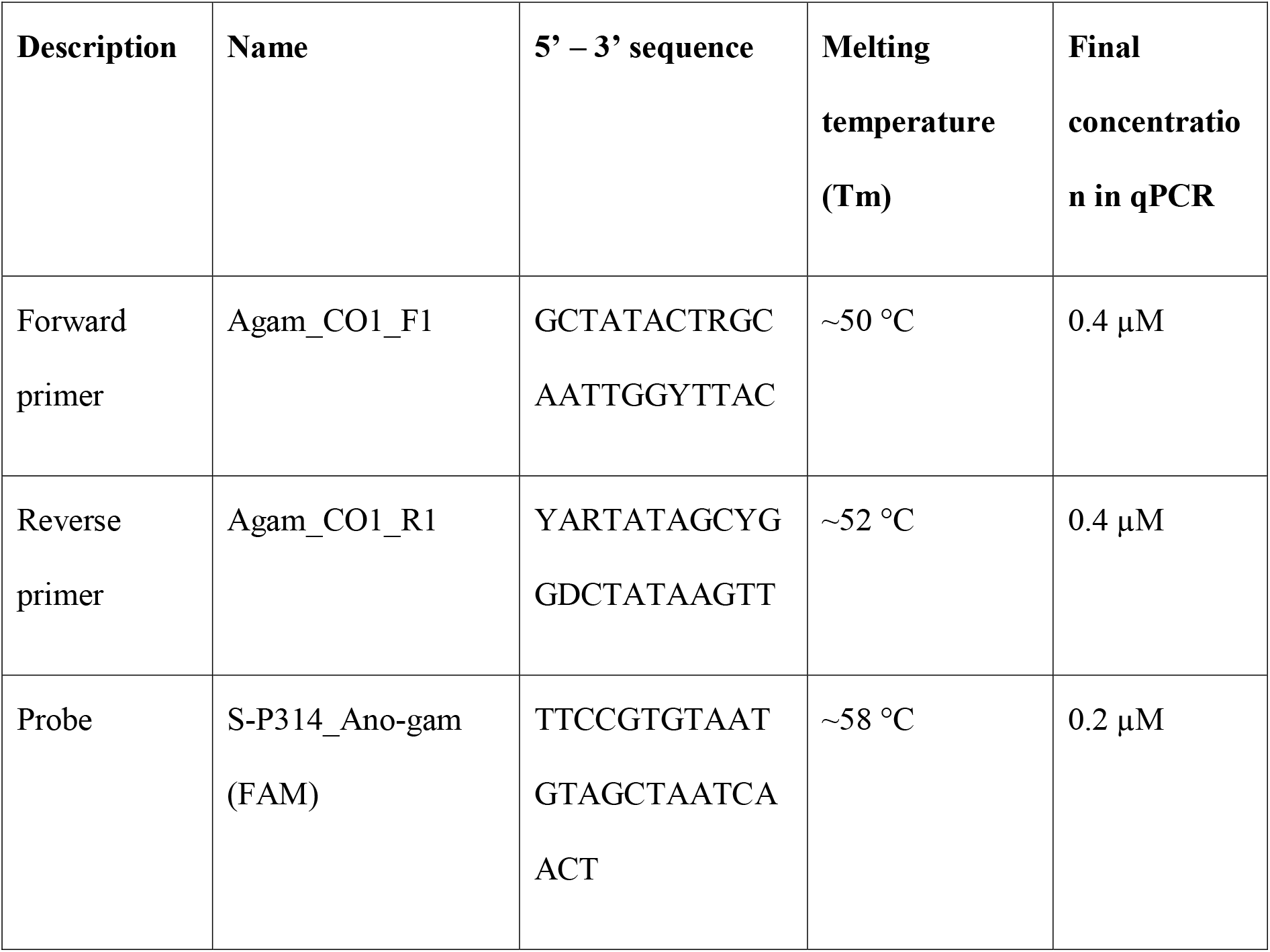
Primers and probe for An. gambiae detection qPCR assay.

### In vitro

The primers and probe were tested using the Luna® Universal Probe qPCR Master Mix (New England Biolabs, #3004L) in a total volume of 10 µl containing 3 µl of template DNA, 1x Luna Universal Probe Mastermix and each primer/probe at its corresponding concentration (Table 1). Amplifications were carried out under the following thermocycling conditions: 1 min at 95 °C, 40 cycles of 15 s at 95 °C (denaturation) and 1 min at 56 °C (annealing/extension).

Several individuals of target and non-target species (target: 3 *An. gambiae* s.s., 2 *An. coluzzii*, 3 morphologically identified *An. gambiae* complex from Angola [33,34], non-target: 1 *Aedes japonicus*, 1 *An. plumbeus*, 1 *An. funestus*, 1 *Culex pipiens*) were tested as original DNA extracts and in dilutions of 1:10 and 1:100 in the qPCR. All mosquito samples (target and non-target species) were transferred from the storage tube (dried or ethanol) or to a new tube containing lysis buffer (200 µl TES and 5 µl Proteinase K) to extract DNA. Samples were then homogenized with glass beads for 2x 20 sec at 5500 rpm (Precellys, Bertin Technologies) and incubated overnight at 58 °C. Total DNA was subsequently extracted using the DNeasy Blood & Tissue Kit (Qiagen) on an automated platform (Kingfisher Flex 96, ThermoFisher Scientific).

The sensitivity of the qPCR assay was tested on serially diluted DNA templates ranging from 1,000 to 2.5 double-stranded (ds) copies µl^-1^ of synthetic DNA template. The synthetic gene was produced by Eurofins Genomics (Ebersberg, Germany) and provided in the form of a plasmid. The 300 bp fragment was retrieved from a target COI sequence, Genbank accession number DQ465336.1. After dissolving the lyophilized DNA and linearising it by restriction digest, the corresponding dilutions of the synthetic gene (ranging from 1,000 to 2.5 double-stranded (ds) copies µl^-1^ of DNA template) were prepared and tested in the sensitivity PCR. Note that the final number of template molecules in the PCR depends on the volume of template DNA used; here, when 3 µl of template DNA of a 10 copies µl^-1^ concentration is used, the actual number of ds template molecules subjected to PCR is 30. Cq values were determined as the cycle number at which baseline-corrected fluorescence (ΔRn) crossed the software-defined threshold positioned within the exponential phase of amplification.

### In silico validation

The primers were used in primer-BLAST [35,36] against a local download of the NCBI nucleotide database [37] to assess amplification of both target and non-target sequences, using an e-value of 1,000 and word-size of 7, outputting all available taxonomic fields of BLAST matches. Sequences were considered a potential match if ≥20 of the 22bp of both primers matched the sequence, in regions 150-300bp apart (giving an amplicon of the desired length). All *in silico* analyses are fully documented at https://github.com/hemprichbennett/qpcr_anopheles_gambiae.

## Results

### In vitro

DNA of the morphologically identified *An. gambiae* complex mosquitoes, plus *An. gambiae* s.s. and *An. coluzzii* (both *An. gambiae* complex members) was successfully amplified with Cq values ranging from 18 to 25 (original extract to 1:100 dilution of target DNA extract from adult mosquitoes, Figure 1), with no other species amplifying. Sensitivity tests showed that the qPCR assay was highly sensitive, successfully amplifying the target DNA down to 5 copies µl^-1^ (i.e. 15 copies per reaction) with Cq values ranging from 29 to 37 (1,000 to 5 copies µl^-1^; Table 2, Figure 2).

**Table 2:** Sensitivity test results of the specific qPCR assay for *An. gambiae*.

| Concentration<br>(copies $\mu\text{l}^{-1}$) | Number<br>of<br>replicates | Number positive | Mean Cq | S.d. Cq |
| --- | --- | --- | --- | --- |
| 1000 | 1 | 1 | 28.75 |  |
| 500 | 1 | 1 | 30.33 |  |
| 100 | 1 | 1 | 33.52 |  |
| 50 | 4 | 4 | 35.08 | 0.53 |
| 25 | 3 | 3 | 35.78 | 0.57 |
| 10 | 4 | 2 | 35.54 | 0.1 |
| 5 | 4 | 1 | 37.41 |  |
| 2.5 | 1 | 0 |  |  |

**Figure 1:**
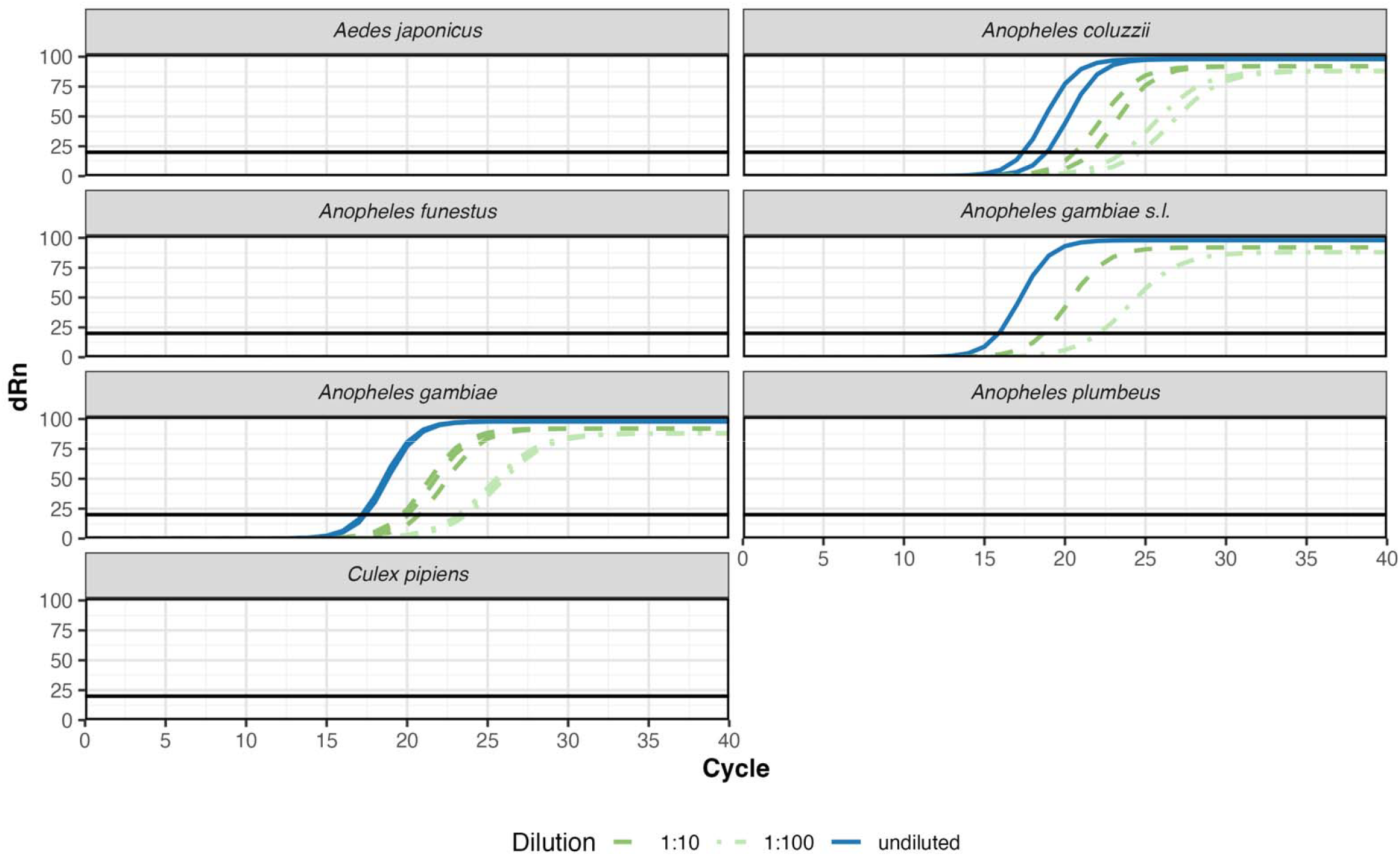
qPCR amplification plot showing the number of PCR cycles before attaining a given level of fluorescence. The black line indicates the software-defined fluorescence threshold used for Cq (quantification cycle) determination. Fluorescence values are baseline-corrected (ΔRn).

**Figure 2:**
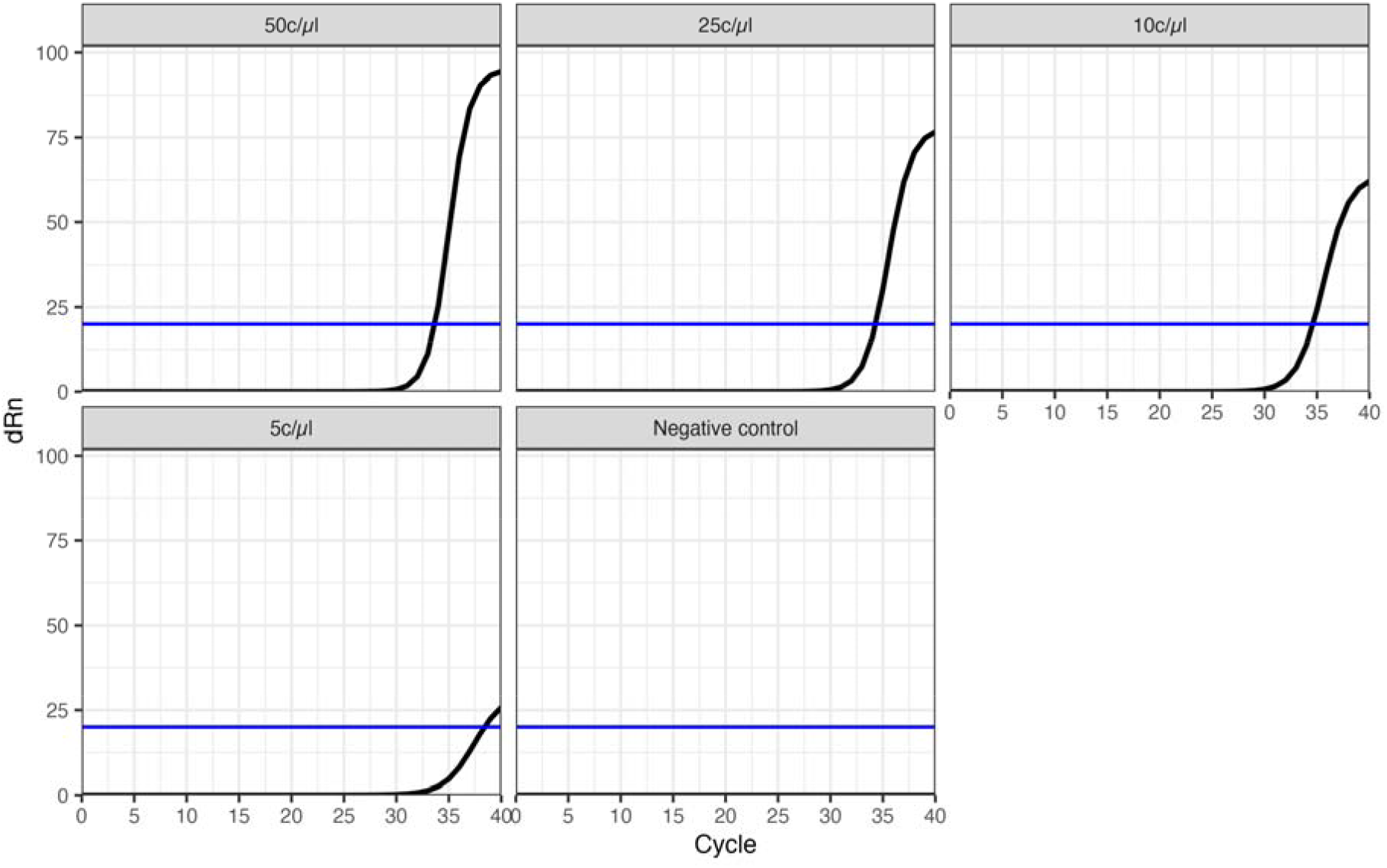
qPCR amplification plot of selected samples of the sensitivity test, showing the initial starting concentration of DNA. The blue line indicates the software-defined fluorescence threshold.

### In silico

No non-target sequences provided a viable amplicon, with no sequences matching the full 22bp of either primer or matching >17 bp of both. The putative ‘amplicons’ of those non-target sequences were >10 million base pairs, so are inviable. The only viable amplicons created were *An. arabiensis, An. coluzzii, An. gambiae* s.s., *An. melas, An. merus* and *An. quadriannulatus*, all members of the *An. gambiae* complex.

## Discussion

The qPCR assay successfully amplified *An. gambiae* complex DNA, showing its utility for detecting prey *An. gambiae* within a predator’s sample. Importantly, the *in silico* testing showed that they did not amplify any non-target sequences from the entirety of Genbank’s nucleotide database, giving high confidence in their specificity. As the primers were shown to be highly sensitive, they can be expected to detect trace amounts of *An. gambiae* within dietary samples, even when the prey has been digested by the predator. The primer amplicon is relatively short at only 192bp long, which is likely to improve its utility in comparison to any candidate primers using longer amplicons, as degradation impedes the detection of longer regions of DNA. The use of a probe in the assay increases the specificity of the detection relative to non-probe-based reactions [38].

Taxonomic and genetic differentiation between members of the *An. gambiae* complex is complicated [39,40], with multiplex PCR assays typically being required to distinguish the complex’s species [26]. Because of the high rate of polymorphisms at all viable binding sites, we opted to use degenerate base pairs in our primers, a practice typically avoided when possible as the primer mixture will contain a lower number of perfectly complementary primer molecules for any given template sequence. However, even with this limitation, our primers proved highly sensitive at detecting low concentrations of *An. gambiae* template. Future studies may improve upon these primers by using genes other than CO1 for analysis, if they are able to find regions that are more conserved within the *An. gambiae* complex and sufficiently different from all other taxa. Our development was restricted to the CO1 gene due to it having far greater availability of both target and non-target CO1 sequences in reference databases than rival genes, but this constraint will be reduced as availability of data for alternative genes increases. As the binding site is located within a mitochondrial gene these primers are expected to encounter higher template DNA availability than primers based on nuclear genes [41], a key consideration when aiming to detect trace levels of DNA.

Our primers are designed to detect very low concentrations of DNA in degraded samples, primarily to identify ecological interactions involving members of the *An. gambiae* complex. Analyses using these primers therefore reliably identify predator taxa that interact with *An. gambiae* but do not indicate dietary preferences or relative dependencies. This is especially true in generalist insectivorous taxa that may consume a wide variety of small Diptera and other small flying insects [42] and with the variable nature of trophic interactions in space and time [43]. Establishing true dietary dependencies would require additional analyses, for instance using DNA metabarcoding or metagenomics to detect other prey that are consumed [22], combined with quantitative network modelling to predict potential shifts in interactions [44].

While these primers were designed for use in dietary study, they may also be of use in other contexts where low levels of *An. gambiae* DNA need to be detected, e.g. in eDNA samples of small water bodies that are potential mosquito breeding sites [28,29], in indiscriminate bulk trap samples (e.g. malaise traps), or detecting whether *An. gambiae* has visited flowers for pollination studies [45]. While the primers only recover single-species information from any of these mixed materials, their greatly improved sensitivity and cost compared to whole-community techniques such as metabarcoding allow them to be used in situations where metabarcoding would not be suitable, or as a complement to it. While medical entomology research often focusses on individual species and subspecies within the *An. gambiae* complex, we chose to target the complex as a whole, as, for ecological research, the loss of taxonomic resolution is outweighed by the value of a single simple assay indicating the presence or absence of a key malaria vector. In studies requiring finer within-complex resolution, the primers described here could act as an initial screening tool to identify samples for further, more specific assays.

## Conclusion

The qPCR primers introduced here will allow advances in the study of predation on *An. gambiae* complex mosquitoes, making it relatively easy and cost-effective to identify if a predator has consumed these important disease vectors. While these primers were designed for use in dietary studies, their utility is equally valid for any studies identifying the presence or absence of these vector species in alternative sources of bulk or degraded material. The addition of these primers to the disease ecology and medical entomology toolkit will enable new insights into the study of malarial mosquitoes, with the ‘simple’ and low-cost nature of the protocol allowing research to take place at scale while complementing other avenues of research.

## Acknowledgements

We thank Sinsoma for their assistance with primer refinement, probe generation and *in vitro* testing. We also thank Gregory C. Lanzaro for contribution of *An. coluzzii* for use in the *in vitro* specificity testing. We thank the Ministry of Health of Angola and the National Malaria Control Programme for their contribution to the research which provided the *An. gambiae* complex samples.

## Availability of data and materials

All data are freely available from NCBI, or in the GitHub repository https://github.com/hemprichbennett/qpcr_anopheles_gambiae.

## Authors’ contributions

DRHB: conceptualisation, methodology, formal analysis, data curation, writing (original draft and review and editing), visualisation

GA: Investigation, contribution of resources, writing (review and editing)

AB: Project administration, writing (review and editing)

FAA: Funding acquisition, supervision, writing (review and editing)

OTL: Funding acquisition, supervision, writing (review and editing)

TDH: Conceptualisation, methodology, supervision, project administration, writing (review and editing)

## Ethics approval and consent to participate

Collection of *An. gambiae* complex samples from Angola was approved by the Instituto Nacional de Investigação em Saúde de Angola (INIS). Reference of the approval: no.12C.E/MINSA.INIS/2022.

## Consent for publication

All authors consent to the publication of this manuscript.

